# *In vivo* validation of an *in situ* calibration bead as a reference for backscatter coefficient calculation

**DOI:** 10.1101/2024.02.07.579320

**Authors:** Yuning Zhao, Gregory J. Czarnota, Trevor H. Park, Rita J. Miller, Michael L. Oelze

**Author notes:** Corresponding Author: Michael L. Oelze, Beckman Institute, 405 N. Mathews, Uni-versity of Illinois at Urbana-Champaign, Urbana, IL 61801; Email.

## Abstract

**Objectives:** The study aims to assess the capability of Quantitative Ultrasound (QUS) based on the backscatter coefficient (BSC) for classifying disease states, such as breast cancer response to neoadjuvant chemotherapy and quantifying fatty liver disease. We evaluate the effectiveness of an *in situ* titanium (Ti) bead as a reference target in calibrating the system and mitigating attenuation and transmission loss effects on BSC estimation.

**Methods:** Traditional BSC estimation methods require external references for calibration, which do not account for ultrasound attenuation or transmission losses through tissues. To address this issue, we use an *in situ* titanium (Ti) bead as a reference target, because it can be used to calibrate the system and mitigate the attenuation and transmission loss effects on estimation of the BSC. The capabilities of the *in situ* calibration approach were assessed by quantifying consistency of BSC estimates from rabbit mammary tumors (N = 21). Specifically, mammary tumors were grown in rabbits and when a tumor reached 1 cm or greater in size, a 2-mm Ti bead was implanted into the tumor as a radiological marker and a calibration source for ultrasound. Three days later, the tumors were scanned with a L-14/5 38 array transducer connected to a SonixOne scanner with and without a slab of pork belly placed on top of the tumors. The pork belly acted as an additional source of attenuation and transmission loss. QUS parameters, specifically effective scatterer diameter (ESD) and effective acoustic concentration (EAC), were calculated using calibration spectra from both an external reference phantom and the Ti bead.

**Results:** For ESD estimation, the 95% confidence interval between measurements with and without the pork belly layer was (6.0,27.4) using the *in situ* bead and (114, 135.1) with the external reference phantom. For EAC estimation, the 95% confidence interval were (−8.1, 0.5) for the bead and (−41.5, −32.2) for the phantom. These results indicate that the *in situ* bead method shows reduced bias in QUS estimates due to intervening tissue losses.

**Conclusions:** The use of an *in situ* Ti bead as a radiological marker not only serves its traditional role but also effectively acts as a calibration target for QUS methods. This approach accounts for attenuation and transmission losses in tissue, resulting in more accurate QUS estimates and offering a promising method for enhanced disease state classification in clinical settings.

## Introduction

Spectral-based quantitative ultrasound (QUS) imaging is a technique that utilizes information from the frequency content of the backscattered signal from tissues to classify tissue state. One spectral-based parameter is the backscatter coefficient (BSC), which can be modeled to relate the frequency spectrum of backscattered ultrasound to the density, size, and shape of scat-terers in the tissues. Differences in scattering structures can be used to differentiate cancer from noncancerous tissues (1), detect changes in scatter-ing structure due to heating (2), characterize liver disease (3), (4), (5). A review of applications for QUS can be found in (6).

Several studies have indicated that cells and cellular structure play an important role in the scattering of ultrasound from tumors. Tumor treatment can alter the mechanical properties and structures of tumor cells, which can lead to changes in tissue backscatter (7). The results of a study by (8) showed a relationship between tumor cell death and the BSC slope extracted from the backscattered signal power spectrum. Both high-frequency (20-50 MHz) and low-frequency (7 MHz) QUS have demonstrated the ability to detect microstructure changes during tumor death (9), (10), (11) and predict the response to neoadjuvant chemotherapy (NAC) (12). These findings suggest that QUS can provide valuable metrics for monitoring tumor treatment.

Properly classifying disease states of tissues using spectral-based metrics like the BSC depends on the ability to accurately and consistently estimate the BSC under different imaging scenarios. For example, it is important to account for system settings, transducer diffraction, attenuation of the ultra-sound signal and transmission loss between multiple layers (13). Although traditional calibration methods, such as the planar reflector (14), (15), (16) or the reference phantom technique (17), can account for the effects of the transducer and system settings, they are limited in their abilities to effec-tively account for the attenuation and transmission losses (18), (19). Several approaches have utilized the total and local attenuation method to attempt to compensate for attenuation losses (19), (20), (21). Coila et al. proposed a full angular compounding method to create maps of attenuation using ultra-sound computed tomography to account for attenuation losses in the estima-ton of the BSC (22). While the clinical feasibility of these techniques shows promise, there remains high variance in estimates of the attenuation. It is also noted that some methods, such as the spectral log difference approach, have been developed to account for transmission losses between layers, albeit with varying degrees of effectiveness.

This effect may be exacerbated in applications such as monitoring changes in tumor structure over time due to neoadjuvant chemotherapy. In this case, ultrasound scans will target the same tissue region at different times, e.g., before onset of chemotherapy, one week post therapy, four weeks post therapy. Perfectly orienting the transducer to acquire the exact same image plane is not possible from one scan to the next. Therefore, there can be different layers and attenuation profiles to a targeted tissue region from one scan to the next. Unless the overlying tissue attenuation and transmission losses are accounted for properly, additional bias and variance of QUS estimates will be introduced leading to poorer classification.

To overcome the transmission and attenuation loss problem, (23) pro-posed an *in situ* reference method. The *in situ* reference, i.e., a small metal bead, involves utilizing a reference that is already contained within the sam-ple that is being characterized. Because the reference material is in the sample medium, it shares similar transmission and attenuation losses with adjacent regions in the sample. Therefore, the *in situ* approach can provide a calibration spectrum that also approximately accounts for transmission and attenuation losses. To obtain more reliable and stable BSC estimates us-ing an *in situ* calibration bead, (13) investigated methods to optimize the calibration spectrum by limiting the number of multiple echoes from the bead and interpolation methods for decreasing sampling error. They demon-strated that consistent calibration spectra from the bead could be obtained through averaging spectra backscattered from across the bead surface even if the bead was larger than the beamwidth. More recently, (24) found that the *in situ* calibration bead could also better account for nonlinear distortion of BSC estimates that may occur during propagation compared to a reference phantom technique.

In this study, we focused on *in vivo* validation of the *in situ* calibra-tion approach by quantifying the frequency-dependent scattering parameters acquired from mammary tumors in multiple rabbits. The *in situ* target con-sisted of a 2-mm diameter titanium (Ti) bead that was implanted into the tumors. The precision of estimates using the *in situ* calibration approach was compared to an external reference phantom approach when scanning tumors with and without a lossy layer placed on top of the tumors.

### Theory

Spectral-based parameterization of ultrasound signals is based on the estimation of the BSC. The BSC is a fundamental property of tissue much like sound speed and density. Therefore, if estimated correctly, the BSC can be operator and system independent. The BSC is defined as the differential backscattered cross section per unit volume (15; 6), expressed as:

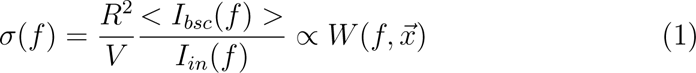

where *σ*(*f*) is the BSC of sample, *R* is the distance to scatter, *V* is the scatter volume, *I_in_* and *I_bsc_* are the incident and backscatter acoustic intensity, and *W*(*f*) is the averaged backscattered power spectrum calculated from the raw radiofreqeuncy (RF) ultrasound backscattered signals. *W*(*f*, 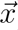) can be modeled as:

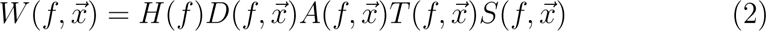

where *H*(*f*) is the system impulse response including excitation voltage, *D*(*f*, 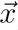) is the transducer diffraction factor and is position dependent, *A*(*f*, 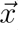) and *T*(*f*, 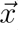) are the attenuation and transmission loss during propagation in tissues, while *S*(*f*, 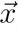) is the scattering function describing the underlying tis-sues.

To isolate the scattering function *S*(*f*, 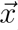) from the averaged backscattered power spectrum *W*(*f*, 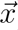), we need to eliminate the other factors. An external reference method, such as the reference phantom method (17), can capture *H*(*f*) and *D*(*f*, 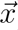) by scanning a separate reference phantom with known parameters using the same transducer and system settings (see Fig. 1):

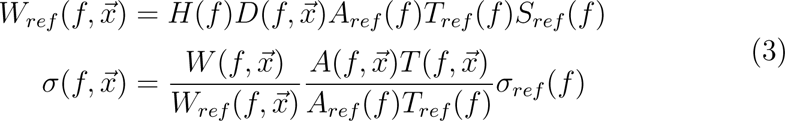

**Figure 1:**
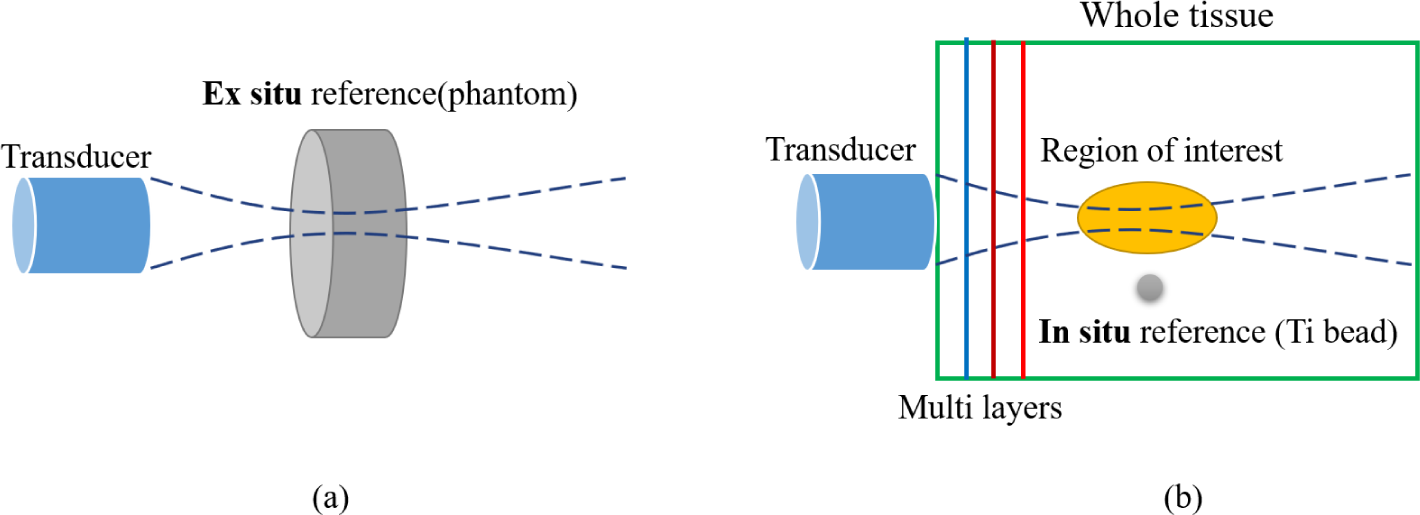
(a) The traditional reference phantom method, i.e., scanning an external phantom with the same transducer as reference and (b) the new *in situ* reference method using a bead at around the same depth as the region of interest.

where *σ_ref_* (*f*) is the known BSC of the reference, *A_ref_* (*f*) and *T_ref_* (*f*) are the attenuation and transmission loss in the reference. The attenuation could be estimated in theory but in experiments with complex structures like multiple tissue layers the accuracy of the attenuation estimates is unknown and the variance of attenuation estimates can be large. Furthermore, due to propagation through multiple tissue layers, transmission loss *T* (*f*) cannot be fully captured by an external reference method.

The *in situ* bead reference assumes that the transmission and attenuation loss of tissue at the location of bead is approximately the same as the transmission and attenuation loss of adjacent tissues at the same depth as the bead. In addition, the backscattered signal from the bead is much larger than the signal from the surrounding tissue. Under these two assumptions, and keeping the bead and tissue scanning system settings the same, the BSC for a sample can be estimated from the backscattered power spectrum calculated from the bead:

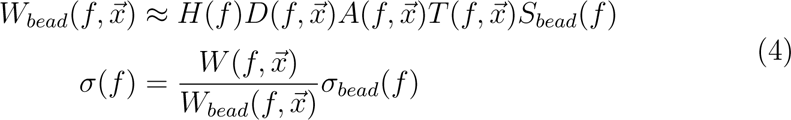

where *W_bead_*(*f*, 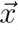) and *σ_bead_*(*f*) are the bead power spectrum and BSC (23). Attenuation and transmission losses in the tissue layers, which are approximately the same for the bead, are eliminated by division. Because the BSC is system and operator independent, the BSC of a bead of particular material and size only needs to be calculated once for subsequent experiments.

## Materials and Methods

### Bead BSC calculation

To use the Ti bead as a reference, one must first estimate the BSC from the Ti bead over the analysis bandwidth. In this subsection, we provide the details on how the BSC from the Ti bead, used as a reference in the rabbit experiments, was estimated. The procedure consisted of creating a phantom with a single bead embedded within the phantom. The same array transducer used in the rabbit experiments was used to insonify the bead and measure the ultrasound backscattered from the bead. Based on the measured backscattered signals, the BSC versus frequency was estimated for the bead.

The BSC of a 2-mm Ti bead was estimated by measuring the ultrasound backscattered signal from a bead embedded in a tissue-mimicking phantom. The phantom consisted of 9g of agar, 3g glass beads (75-90*µm* in diam-eter), and 300mL degassed, deionized water. The mixture was heated to approximately 80°C. To support the Ti bead, half of the mixture was placed in a pre-cooled container for one minute, after which the bead was added, followed by the remaining mixture. The phantom was then placed in a ro-tisserie, which rotated the phantom continuously for 30 minutes to prevent the bead from sinking to the bottom as the phantom hardened.

The ultrasound backscatter was measured from the bead in the phantom using an L14-5/38 probe connected to a SonixOne ultrasound imaging scan-ner (BK Ultrasound, Peabody, MA), which was also used in the *in vivo* tumor experiments. B-mode images of the bead embedded phantom were displayed and recorded using the SonixOne machine, while the L14-5/38 probe was po-sitioned using a LabVIEW-controlled micropositioning system (Daedal, Inc., Harrison City, PA). To calculate the BSC of the bead using equation (3), both the power spectrum of the bead and the power spectrum of the phan-tom at the same depth were estimated from ultrasound backscatter. The probe was swept twice across the phantom to acquire multiple scan lines of the phantom for later compounding.

### In vivo Data Collection

The protocol for the described experiment was approved by the University of Illinois at Urbana-Champaign Institutional Animal Care and Use Com-mittee (IACUC protocol 20087). In total, 21 separate rabbit tumors were scanned in this study. A single rabbit experiment spanned from 3-5 weeks and comprised three stages:

1. Tumor injection and growth: VX2 tumor fragments were injected into the mammary fat pad of a 2 kg female New Zealand White Rabbit, under anesthesia using 2% isoflurane. The tumor growth was monitored for the following 2-3 weeks until the tumor diameter reached around 2.5 cm.
2. Bead injection: When the tumor reached 2.5 cm in diameter, a 2-mm Ti bead was implanted into the tumor area, with a 12-gauge stainless steel needle after shaving and disinfecting the area.
3. Imaging and data collection: Both on the injection day and three days after the bead injection, the tumor with the embedded bead was imaged using a SonixOne ultrasound imaging scanner equipped with an L14-5/38 transducer. Images were captured with and without a sample of pork belly, around 1.5-2 cm thick, purchased from a local supermarket and placed on top of the tumor to simulate a fat layer. The transducer was scanned across the tumor by free hand translation. This resulted in a 3D volumetric scan of the tumor through stacking of individual frames of the tumor. RF data and corresponding video files were recorded via the SonixOne ultrasound machine. Only data acquired at day three were processed to remove any effects from blood pooling at the injection site on the day of injection, i.e., day 0.

### Bead segmentation

In order to use the bead as an *in situ* reference, imaging frames con-taining the bead and the actual bead location in the imaging frames must be detected for processing the bead signal. Manually selecting the bead location from many image frames is a laborious and challenging task, requiring indi-viduals to scan through all the frames, identify a possible frame, and label the bead location. Automated bead separation can significantly streamline the entire BSC data processing pipeline. An example of the bead segmentation is illustrated in Fig 2.

**Figure 2:**
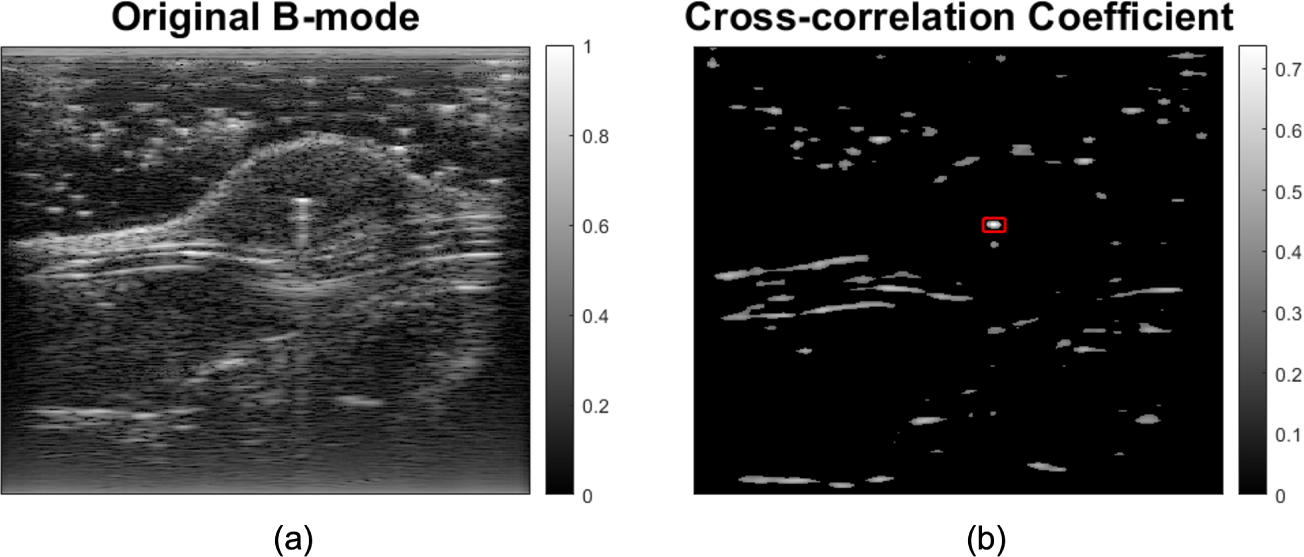
(a) Original B-mode image (b) Cross correlation coefficient with a 0.3 threshold. Max coefficient location is in the red box.

Automatic detection of the bead in the images was accomplished by cor-relating the bead locations in frames containing beads with a reference bead signal. During our scanning process, as the transducer is scanned across the tumor, both the shape of the tumor and the position of the bead can change. This variation necessitates the application of cross-correlation for each individual frame to accurately track the bead’s shifting location.First, a bead scattering function (BSF), which was extracted from six sets of labeled bead data (shown in Fig 3), was selected as a reference. In Fig. 3, the BSF appears similar no matter if additional attenuation is present due to a pork belly layer. This consistency of the BSF suggests that simple tools could be used to detect the presence of a bead in an image frame given sufficient signal-to-noise ratio (SNR).

**Figure 3:**
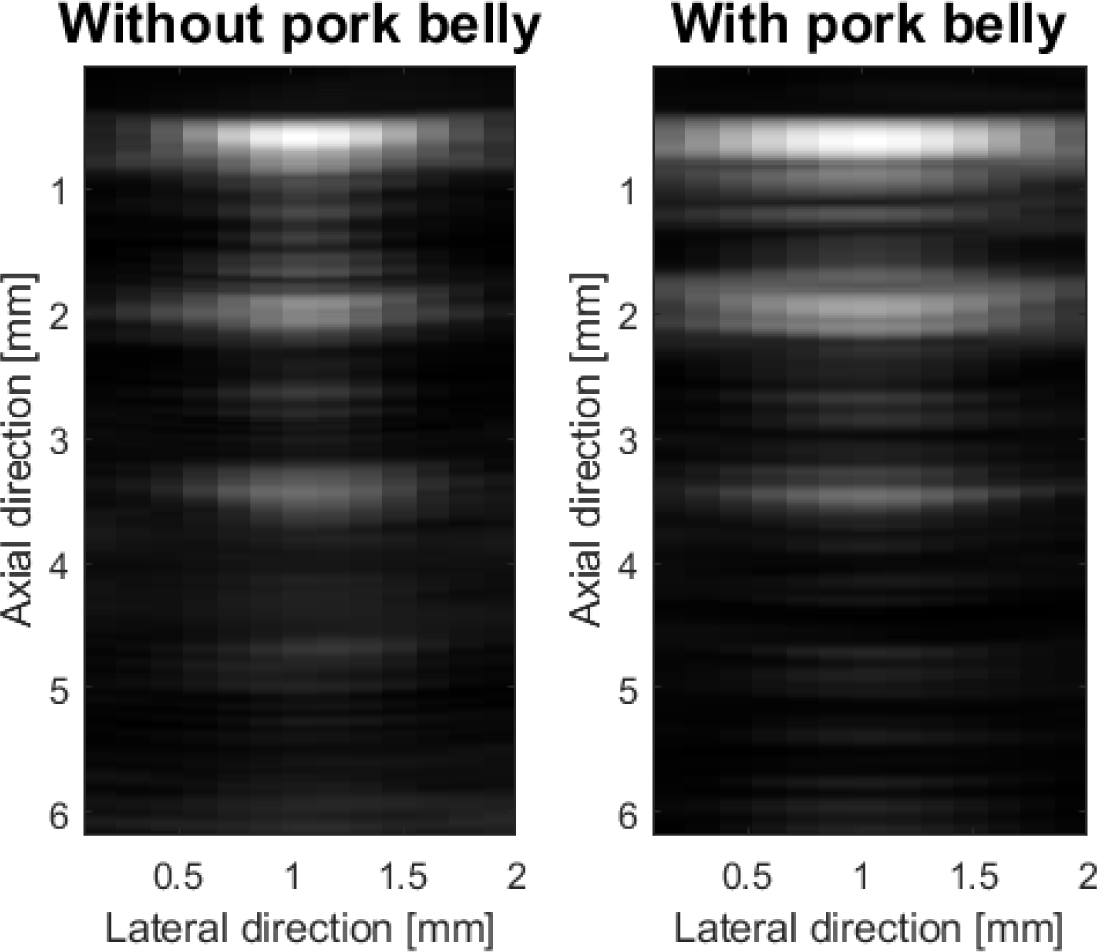
The BSF of the bead shown in B-mode images.

The BSF for the bead includes reverberation from multiple echoes and is consistent between cases where more attenuation of the signal occurs, i.e., when a pork belly layer is on top of the sample. Next, the RF data for different imaging frames were converted to B-mode images using a decibel (dB) scale with a dynamic range of 60 dB. The B-mode images were processed by cross-correlating each frame with the BSF. The maximum coefficient value and its location were recorded for each frame, and this process was repeated for all frames in a tumor data set. Finally, the frame with the maximum cross-correlation coefficient was labeled as the desired frame for calculating the bead reference signal.

The capability to account for attenuation loss in the estimation of the BSC *in vivo* is demonstrated by comparing two distinct conditions: the presence and absence of pork belly. A challenge in this comparison arises from the difficulty in matching exact imaging frames between pork belly and no pork belly conditions. It is virtually impossible to capture the exact tumor volume from two separate scans, especially when pork belly layers are added on top of the tumor. However, the bead signal is not conditioned to whether there is pork belly or no pork belly on top of the tumor because the bead is a sphere and the angle of insonification does not change the backscatter from the bead. The key is to scan across the bead in the tumor so that the maximum bead signal is captured in an imaging frame, which will later be identified and used for the reference.

### Tumor BSC Calculation

For each tumor scan, ten frames were selected for BSC calculation. The ten frames were evenly spaced across the tumor, representing a volume of the tumor, by taking ten frames from the video file of data collected by moving the transducer across the tumor. The center of the ten frames was located at the frame where the maximum bead signal was confirmed. On each side of the bead frame, additional frames were selected (five on one side and four frames on the other side of the bead frame) at even spacing between frames to provide independent frames of the tumor volume. The exact spatial separation between these frames varied because the sizes of the tumors varied. The spacing between frames depended on the distance between the bead and the edge of the tumor on both sides of the tumor in the elevational direction. The frame spacing varied due to the differences in the extent of the tumor captured in each scan, and the spacing between the selected frames ranged between 2 and 16 frames. By adjusting the distance between frames, we were able to capture representative estimates of the whole tumor. For each frame, the tumor region was segmented and then separated into small data blocks (3mm * 3mm, 3 mm is approximately 10 wavelengths of the center frequency (5.5 MHz) of the analysis bandwidth) with 75% overlap and the average BSC of each tumor region was calculated using both the bead reference and reference phantom for comparison. Larger data block sizes could be used to reduce variance but at a trade off in spatial resolution of the ESD and EAC maps (25). The reference phantom (CIRS, Sun Nuclear, Norfolk, VA) had known attenuation (0.67dB/cm/MHz), sound speed (1537 m/s) and BSC measured with two single-element transducers (5 and 10 MHz center frequencies), shown in Fig 4. The phantom had a thin saran layer to act as an acoustic window. The BSC was averaged from the selected ten frames and compared using different methods both with and without pork belly on top of the tumor.

**Figure 4:**
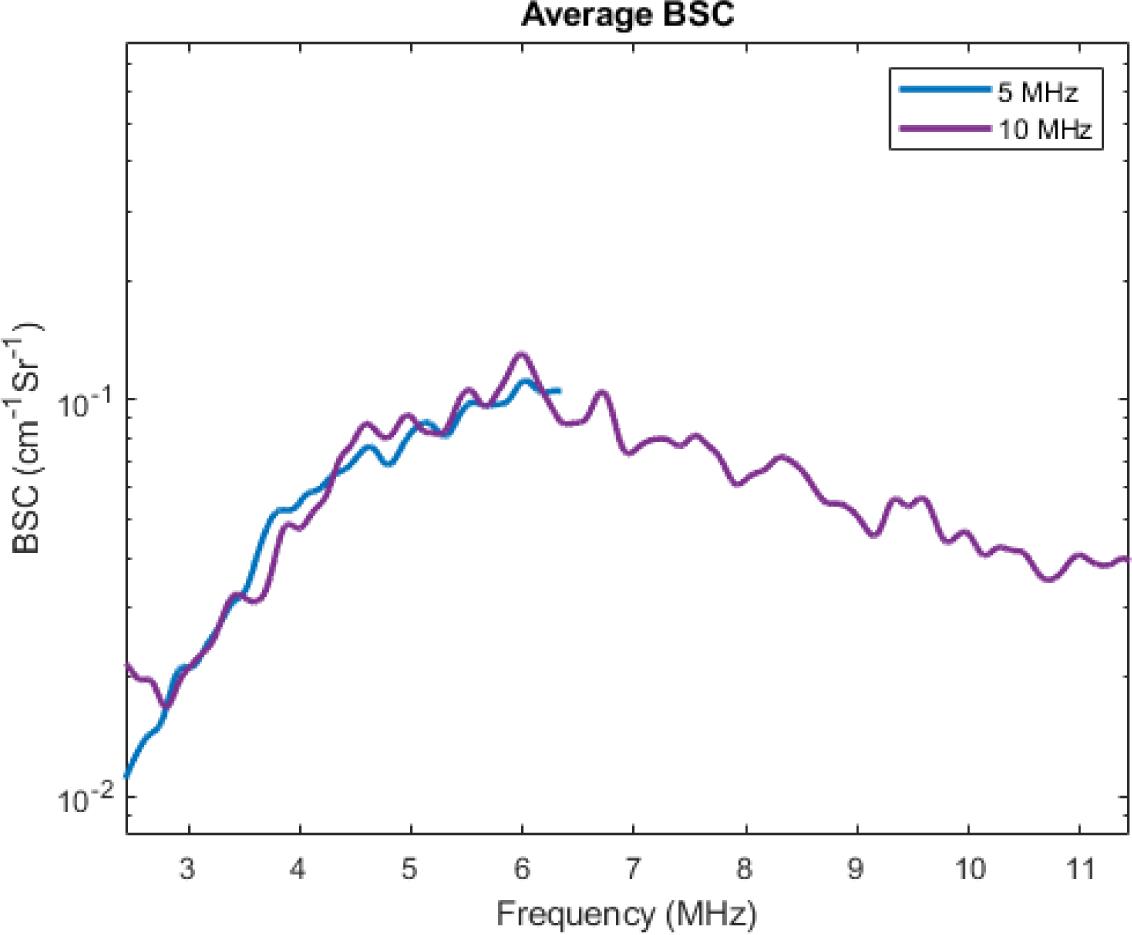
The BSC of the CIRS phantom used in the BSC measurement.

To assess the ability of the *in situ* bead to provide accurate and precise estimates of tumor scatterer properties when different loss scenarios occur between the tumor and transducer, the BSC was parameterized and the resulting parameters tested to determine if statistically significant differences were present. Statistically significant differences would suggest that the calibration approach, i.e., external reference phantom or *in situ* calibration bead, was not able to account for changes in the transmission and attenuation losses to the tumor sample. Two scatterer-related parameters, effective scatterer diameter (ESD) and effective acoustic concentration (EAC), were estimated from the BSC using a Gaussian scattering model (15) (26),

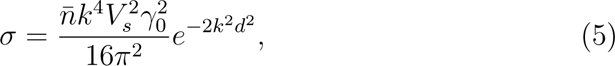

where *σ* is the BSC, *n̅* is the average number density of particles, *k* is the wave number, *d* is the characteristic dimension, *V_s_* = (2*πd*^2^)^3^ is the average particle volume, and *γ*_0_ = (*Z* − *Z*_0_)/*Z*_0_ is the fractional change in impedance between the scatterers and the surrounding medium. For *γ*_0_, *Z* and *Z*_0_ are the scatterer impedance and the surrounding medium impedance.

The relationship between ESD and characteristic dimension *d* is: *ESD* = 3.11d, and the product of *n̅* and 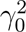 is called effective acoustic concentration (*EAC*). The estimates ESD and EAC provide a physical interpretation of the tumor microstructure and offers features for comparison between tumors when scanned with and without lossy layers placed on top.

### Power spectrum averaging

For all the data collected from the experiments, the power spectrum from the bead needed to be calculated. Only the first echo of the bead was win-dowed, for the purpose of smoothing (13). Next, interpolation and averaging were performed in both lateral and elevational directions because only finite locations were sampled, and the max bead signal location might be missed during the location sampling, shown in Figure 5 (a). The data were interpo-lated via MATLAB (The MathWorks, Natick, MA) using the interp2() func-tion which performs the cubic interpolation, and only the power spectrum from scan lines within the −6 dB rolloff from the bead, as the beam crossed it laterally, were chosen for averaging (13). To quantify the agreement between BSCs estimated with and without a pork belly layer, the root mean squared error (RMSE) was calculated using the bead reference or external reference.

**Figure 5:**
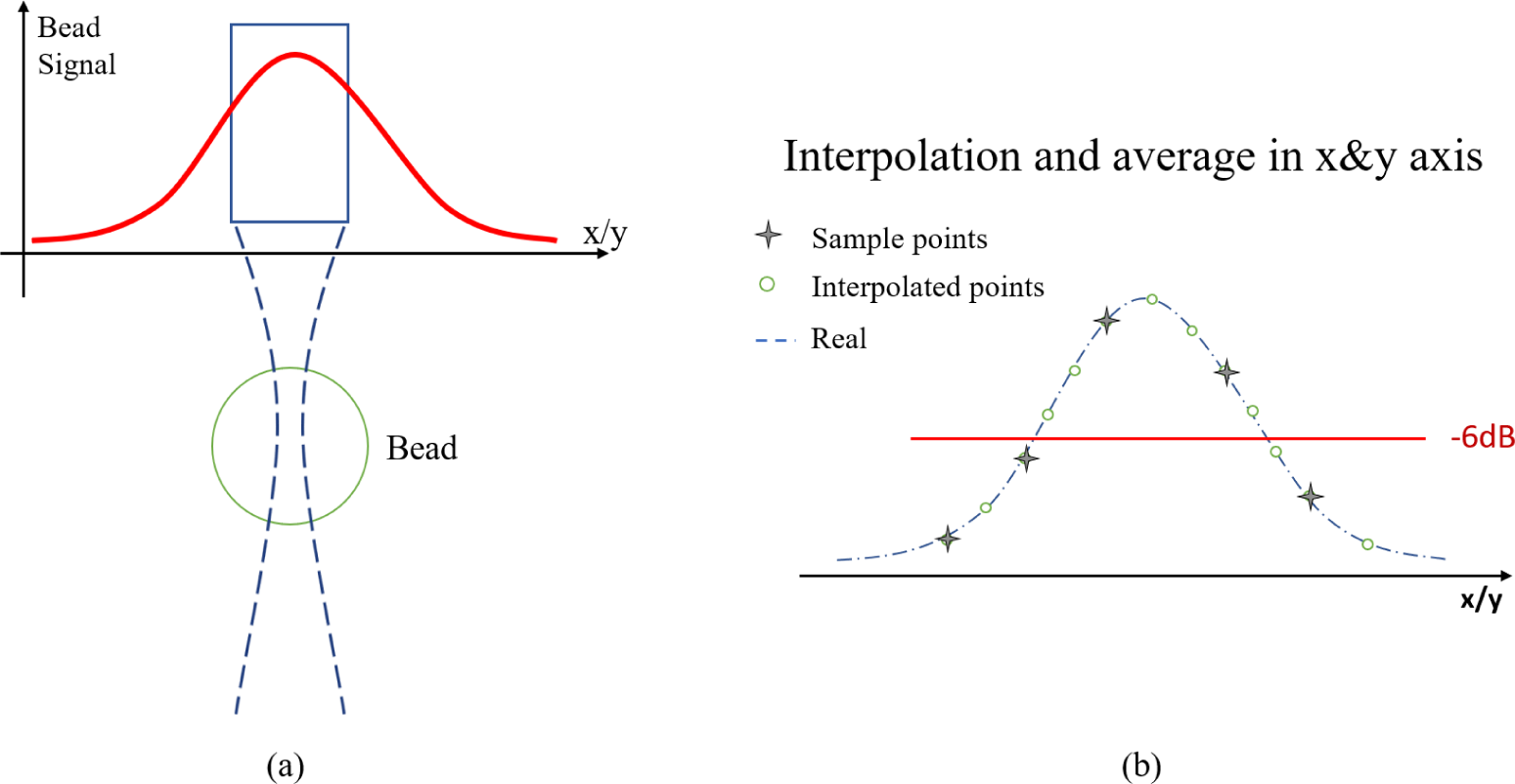
(a) Signal energy as a function of relative location from bead, decreasing from center to edge (b) Example of how the maximum signal off the top of the bead was missed due to finite lateral sampling and signals recovered by interpolation and averaging.

### Statistical analysis

To directly compare the *in situ* bead and reference phantom calibration approaches, a measure of calibration error was defined and applied to each approach separately, then statistically analyzed for overall differences. The measure chosen was the absolute difference in the calibrated QUS parameter (average ESD or average EAC) between two scans of the same tumor: one through the intervening slab (pork belly), and the other without it. This measure serves as a proxy for the “true” absolute error, that is, the error that remains even after calibration and is due to all causes. Though the calibrated parameter value calculated from the scan without the pork belly is not the “true” QUS parameter value, it is presumably closer to the “true” value than the calibrated value calculated from the scan through the pork belly.

A collection of tumors scanned both with and without the pork belly, and to which both calibration approaches had been successfully applied, was selected for this analysis. Because the approaches were applied to the same collection of tumors, a paired analysis was conducted, with each tumor defining a pair. The overall statistical difference in the absolute calibration error was assessed using a paired Wilcoxon signed rank test.

Calibration errors can be attributed to some combination of calibration bias and variance. The bias is the systematic component of the error, possibly positive or negative, that is the same for every instance (tumor). The variance represents the leftover variability (from one tumor to another), after bias is accounted for. To measure bias separately for each calibration approach, a confidence interval for a measure of center called the pseudo-median (associated with a Wilcoxon analysis) can be applied to the pork belly versus no pork belly differences. An interval containing zero would indicate no evidence of bias. To compare variance between the two calibration approaches, a likelihood ratio test can be performed for the null hypothesis of equal variances, using a Wishart model for the covariance matrices of the pork belly versus no pork belly differences. Using such a multivariate approach accounts for pairing by tumor.

## Results

The bead detection method utilizing the BSF and cross-correlation was effective at identifying frames with a bead. In each tumor scan, the BSF correlation technique labeled a frame with the highest cross-correlation co-efficient as the bead frame. Out of 199 tumor image frames, it correctly identified the bead in 174 scans (true positive). However, there were 25 in-stances where it incorrectly labeled a frame (false positive). Consequently, the technique achieved a bead detection rate of 87.4% and a misidentification rate of 12.6%. Given the inherent characteristics of certain tumor tissues, which can resemble the bead and produce robust signals, it’s understandable that these tissues might occasionally be misidentified as the bead. Moreover, in some scans, the bead signals are weak, exhibiting a low SNR, which could also lead to mislabeling.

Figure 6 shows the results of the BSC calculation for a tumor experiment, using both the bead method and the traditional reference phantom method. The RMSE were calculated for BSCs estimated from tumors with the pork belly present and absent for both the *in situ* calibration method and reference phantom method. In our study, which involved 21 tumor scans on the third day after bead injection (Day 3), we carefully analyzed the RMSE values for each scan to evaluate the accuracy of BSC estimation methods. We em-ployed both the bead method and the reference phantom method to calculate RMSEs. In our graphical representation, we observed that the bead method typically demonstrated less variability in RMSEs compared to the phantom method, although there are several exceptions. The RMSE results of these 21 tumors are shown in Figure 7. The averaged RMSE for the *in situ* bead method was 2.7×10^−4^ and for the reference phantom method was 9.3×10^−4^. This suggests that the multilayer loss, including attenuation and transmission, were more properly accounted for using the *in situ* bead method. It should be noted that it is impossible to image at the exact same region in the tumor when scanning with and without meat on top of the tumor, and the tumor tissue is not homogeneous. Therefore, the BSC with and without meat would never match perfectly.

**Figure 6:**
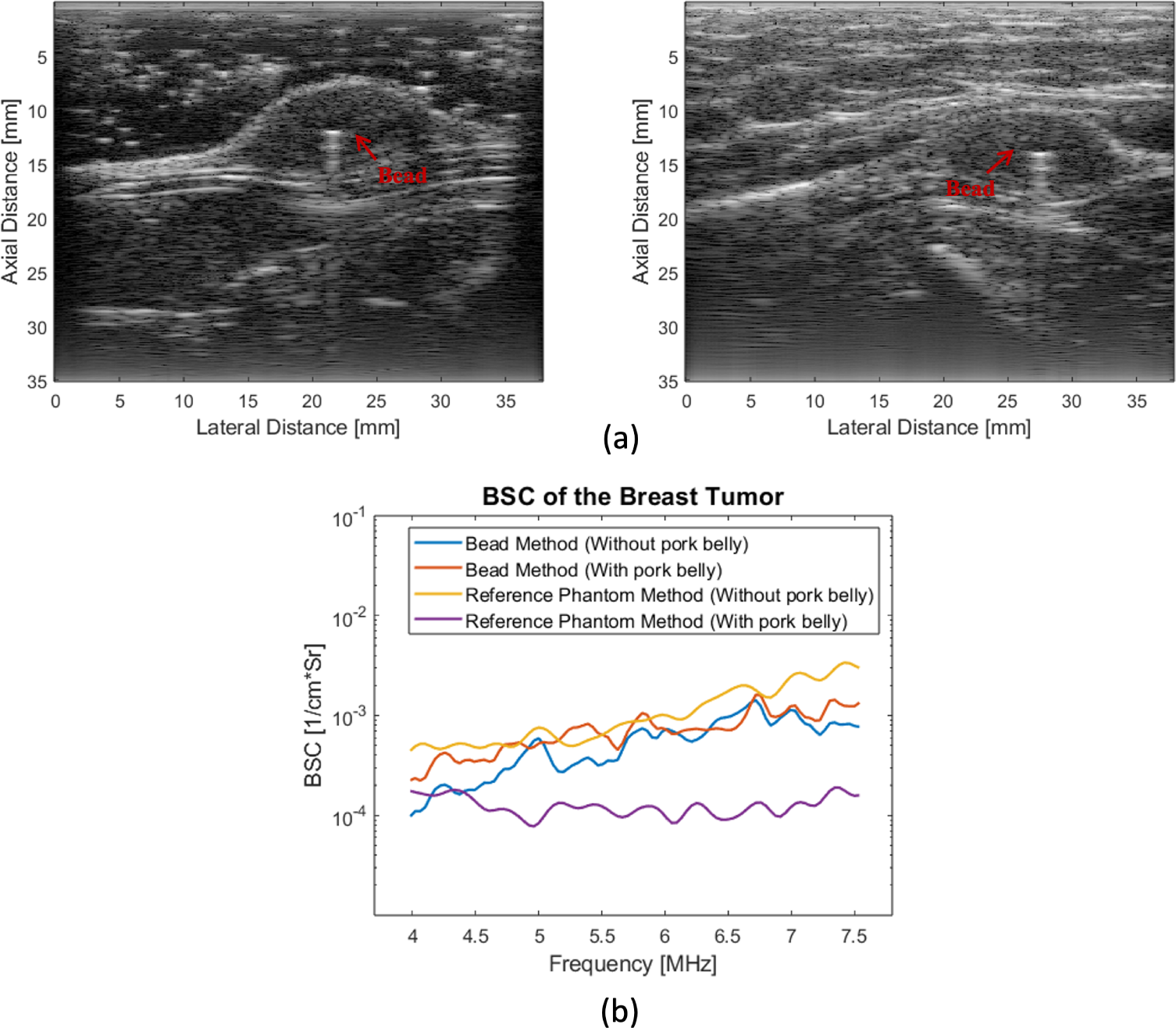
(a) B-mode images of tumor with (right) or without (left) pork belly (b) Corresponding averaged tumor BSC calculated using both the bead method and the reference method.

**Figure 7:**
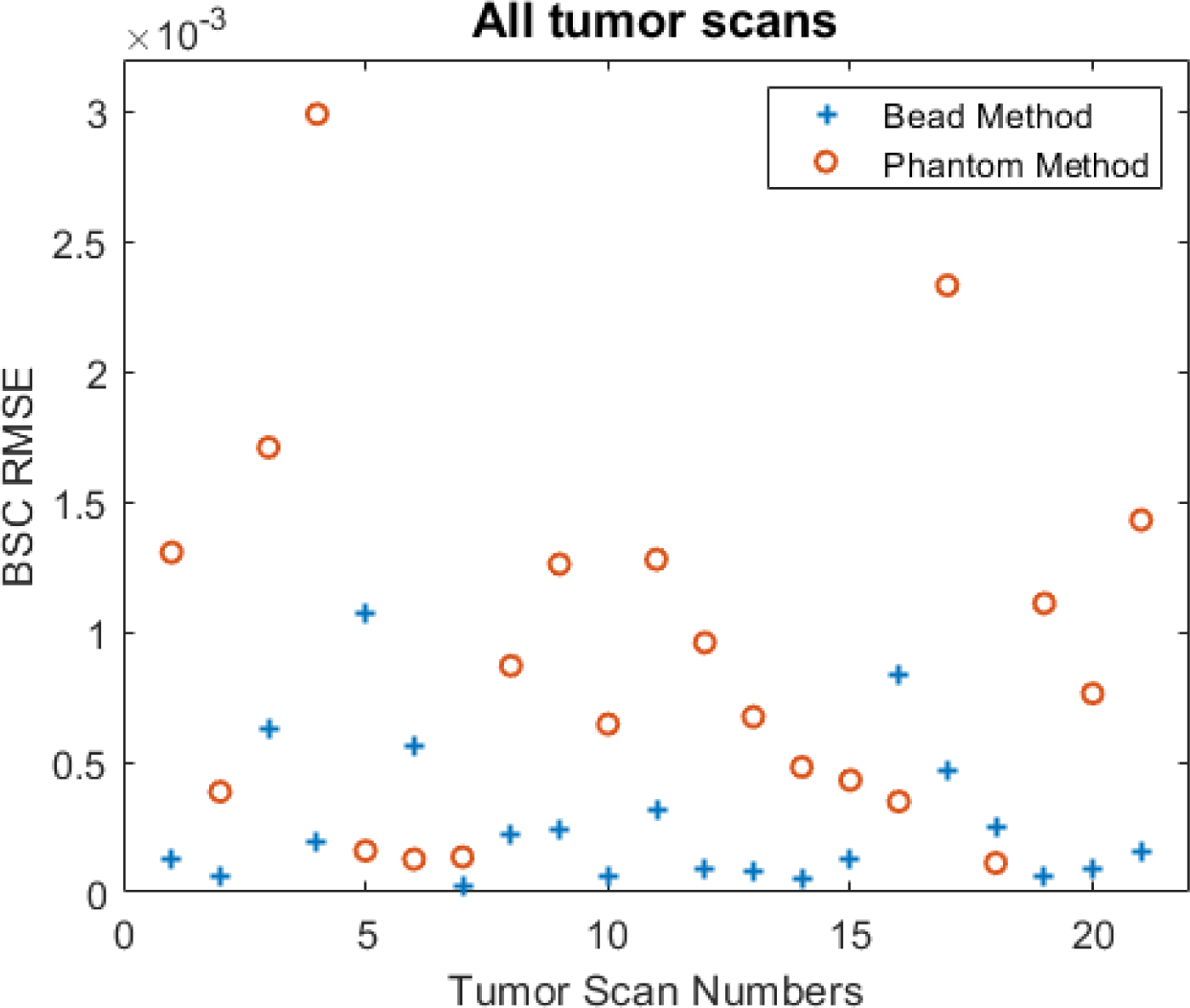
RMSE distances between the Day 3 tumor BSCs from no pork belly and with pork belly cases.

A further investigation was conducted on four cases where the bead RMSE was larger than the RMSE using the reference phantom. We estimated the tumor SNRs of these cases and the the SNR was 10 dB or lower: 6.22dB, 5.81dB, 8.50dB and 10.54 dB. These suggest that the performance of the bead method is closely tied to the tumor SNR, with higher SNR correlating with more reliable outcomes.

Figure 8 presents sample ESD maps of the tumor determined using both the bead and phantom references. With only ultrasound gel on top the tumor, the ESD maps from both methods were comparable due to the gel’s minimal attenuation loss. However, when pork belly was positioned over the tumor, there was a noticeable discrepancy. In the case shown, the ESD values estimated with the phantom reference rose significantly compared to those without the pork belly. In contrast, the *in situ* bead reference method maintained greater consistency in ESD estimation. This indicates that the *in situ* bead reference offers a more reliable BSC calculation by better accounting for attenuation and transmission losses.

**Figure 8:**
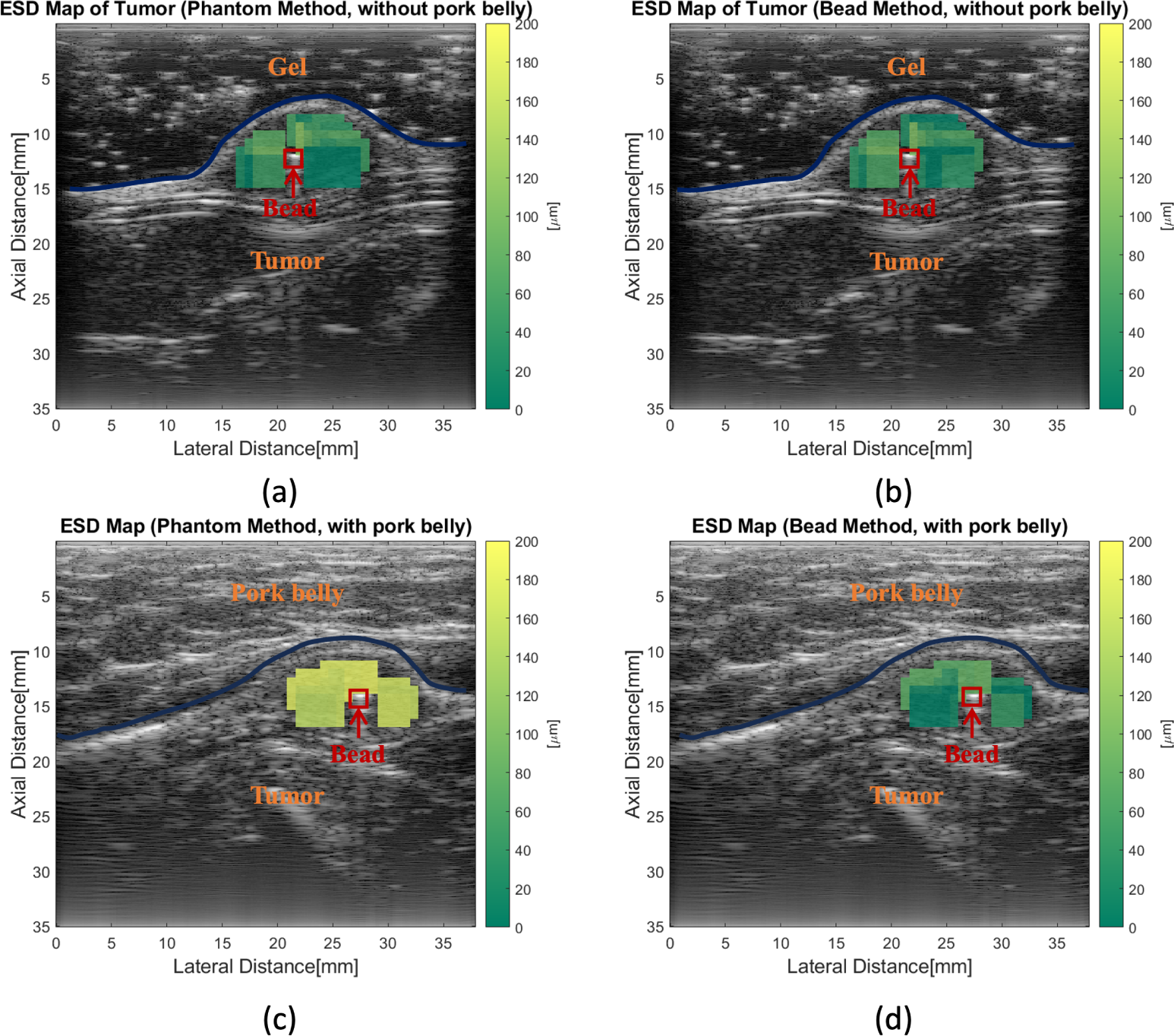
ESD Maps of the tumor using different references in different situations (a) Phantom reference, gel on the top (b) Bead reference, gel on the top (c) Phantom reference, pork belly on the top (d) Bead reference, pork belly on the top.

The measure of calibration error (the absolute difference in the calibrated value between the scan of a tumor beneath the pork belly and the scan of the same tumor without the pork belly) was computed for each calibration method (bead and phantom) and each QUS parameter (average ESD and average EAC). The results appear in the parallel axis plots of Fig 9, which includes only scans acquired on the third day after bead injection (Day 3). Each line segment in the plot represents a different tumor, supporting a paired comparison. As the figures show, for almost all tumors, calibration error was markedly higher for the reference phantom calibration compared to the *in situ* bead calibration, both for average ESD and for average EAC. Paired Wilcoxon signed rank tests strongly support this interpretation, with two-sided p-values of 0.00000024 for ESD and 0.00000047 for EAC (the sam-ple size was *n* = 21 pairs for both ESD and EAC).

**Figure 9:**
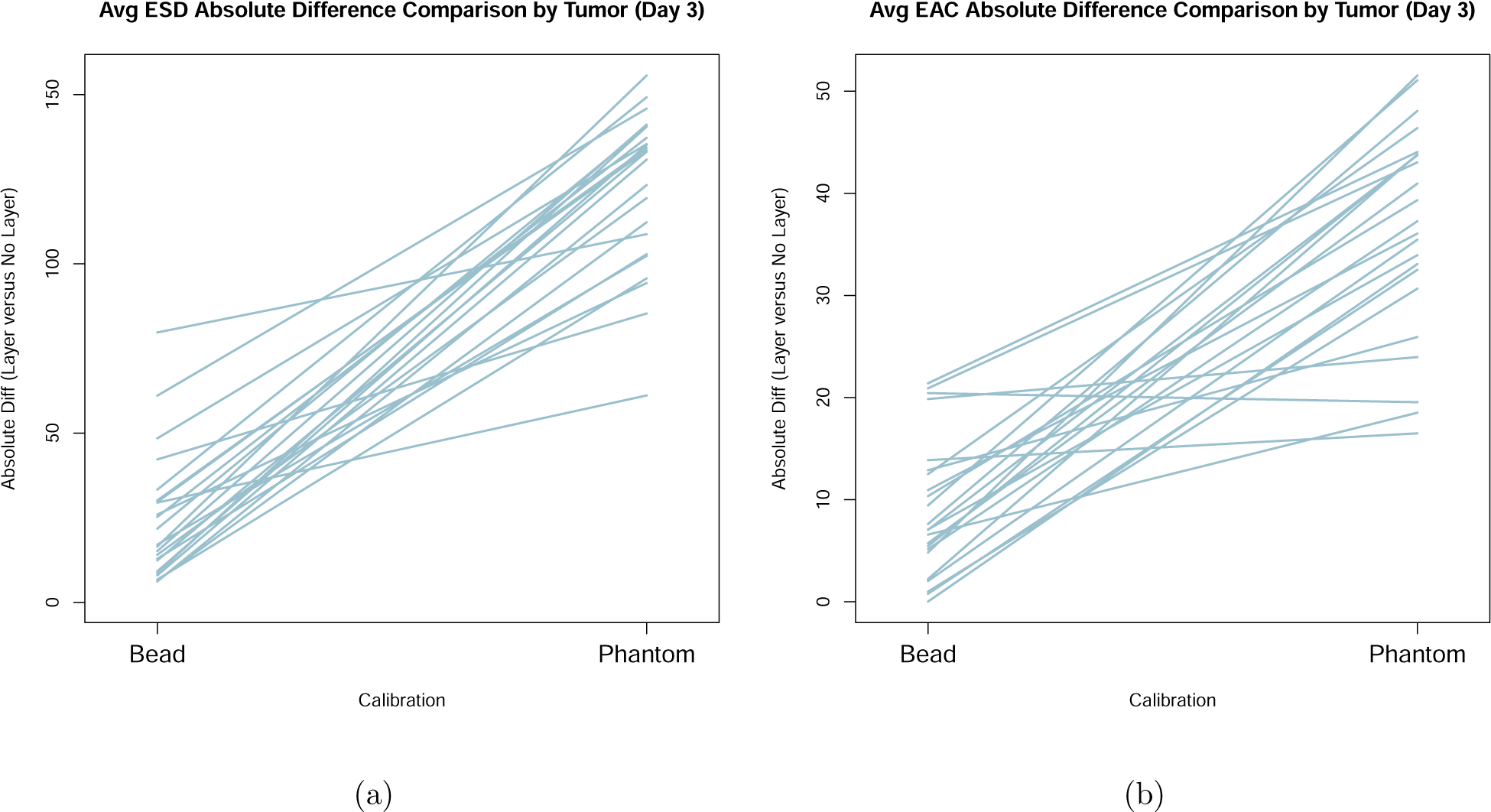
Parallel axis plots of absolute difference calibration error for (a) av-erage ESD and (b) average EAC. Each line is for a different tumor.

Exploration of the separate contributions of calibration bias and vari-ance was based on differences (by tumor) in each QUS parameter between the pork belly and no pork belly scan conditions. For the average ESD, a 95% confidence interval for the pseudomedian (as in a one-sample Wilcoxon analysis) of the differences was (6.0, 27.4) for *in situ* bead calibration and (114.5, 135.1) for reference phantom calibration, indicating that adding the pork belly induced a positive bias for both calibrations, but much higher for the reference phantom method than for the *in situ* bead reference. For average EAC, the intervals were (*−*8.1, 0.5) and (*−*41.5, *−*32.2), indicating a strongly negative slab-induced bias for reference phantom calibration, but not for the *in situ* bead reference. On the other hand, calibration variance was not significantly different between the *in situ* bead reference and the ref-erence phantom method (p-values 0.755 for ESD and 0.997 for EAC, based on marginal likelihood ratio tests that accounted for pairing).

Corresponding analyses conducted for scans acquired on the injection day (Day 0) yielded comparable results and similar general conclusions.

## Discussion

In this study, we conducted the first *in vivo* evaluation of using an *in situ* bead as a calibration source for the estimation of the BSC and related scat-terer properties. The hypothesized benefits of using an *in situ* calibration bead as a reference is the ability to better account for attenuation and trans-mission losses encountered by ultrasound as it propagates through tissue to a region of interest. The performance of the new approach was evaluated in 21 sets of VX2 tumors grown in the mammary fat pad of rabbits to mimic breast tumors. The precision for scatterer property estimates using the bead method was evaluated for multiple tumors.

Based on the comparison and statistical analysis of both ESD and EAC, one can conclude that the bead method is effective in compensating for at-tenuation and transmission losses and potentially nonlinear distortion (24) when compared to the traditional reference phantom method, which used a reference external to the sample. By the Wilcoxon matched-pairs signed-ranks test, the error of the phantom calibration method was mainly due to its much greater bias, rather than to having greater variance. In several samples, however, the estimated average ESD and EAC values had large variations be-tween meat and no meat even using the bead method. This could be caused by the relatively low SNR in tumors, which is a result of high attenuation from the thick and fatty pork belly used in the model. Figure 10 presents an analysis of six cases exhibiting the highest ΔESD between tumors with and without a meat overlay alongside another six cases with low ΔESD val-ues when using the bead reference. From this comparison, we can observe that an SNR for tumors with meat overlays of less than 10 dB were asso-ciated with ΔESD values larger than 40 µm for the bead method. Smaller ΔESD values were observed for larger SNR values in general. This finding underscores the importance of SNR for maintaining good estimates and that, similar to the reference phantom method, the bead method does not account for low SNR situations. This makes sense because the error of the reference phantom method came mainly from the attenuation and transmission loss of the pork belly, which would increase the bias but not influence the variance. The BSC curve of the tumor obtained using the *in situ* bead method and the reference phantom method generally matched well when scanned with-out a piece of pork belly, although the BSC did not completely align. This discrepancy could be due to the attenuation loss compensation factor from the reference phantom method. Furthermore, conducting a longitudinal ex-periment to observe the changes in BSC during tumor growth would yield valuable evidence for the potential clinical application of this *in vivo* bead method.

In the present study, a bead search method was employed to automatically identify the bead in a set of image frames. The success of the identification algorithm will allow a reduction in the time required by a human observer to identify an image frame containing a bead to be used for calibration. The cross-correlation method was utilized for this purpose, which is constrained by the standard bead BSF. The technique worked well under the conditions used in the study with a precision rate of 87.4%. For the failure situations, we manually searched all frames and found the reference frame. Importantly, the peak bead signal was markedly distinct within the tumor images, making it relatively straightforward for human observers to isolate and identify. While manual identification is reliable and ensures that all correct bead frames and locations are found, the automated bead search process offers a significant time-saving advantage. However, varying imaging systems, probes, exper-imental settings, and imaging conditions may necessitate the utilization of different bead BSFs. This is a limitation of the bead segmentation method. Alternatively, computer vision (CV) and other machine learning techniques could be applied for bead detection and warrant further investigation.

**Figure 10:**
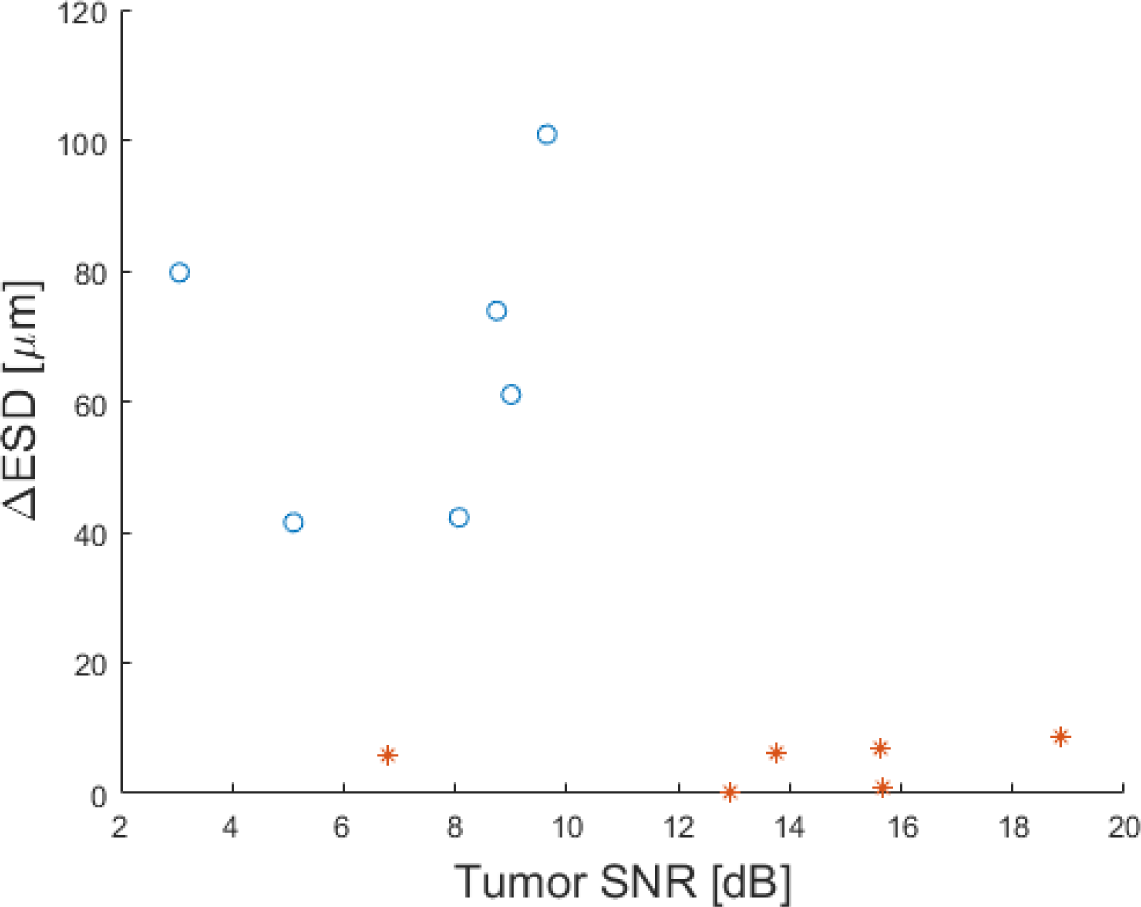
Comparative analysis of ESD and SNR: investigating six scans with high ESD variability (indicated by Blue Circles) versus six scans with low ESD variability (indicated by Red Stars)

In practical terms, the insertion of a small metal bead into a human patient does not face barriers. Currently, radiological clips or makers are used in cancer imaging and during therapy to mark specific tissues, tumors, and lymph nodes. These clips consist typically of metal and come in a variety of shapes and sizes (27). In fact, gold metal beads of 2 mm diameter are already FDA approved for use in patients as a radiological marker, indicating its safety and potential applicability in diverse tissues. For data block size, we chose a 3 mm x 3 mm dimension for BSC estimation, which corresponds to 10 x 10 wavelengths, to balance the variance and resolution of ESD and EAC estimates. While the ideal bead placement is at the tumor’s center to align with its attenuation properties, our findings demonstrate that placement near the tumor’s surface or bottom is also effective, suggesting a flexibility in our method. This flexibility is crucial as it allows for adaptation to the varying shapes and attenuation characteristics of different tumors. However, the closer the region to be interrogated is to the bead location, the better the bead will act as a reference that can account for the attenuation and transmission losses. The use of a spherical metal marker also provides a large impedance contrast with tissue and provides the same backscattered signal no matter the orientation of the transducer probe from the bead.

The success of the *in situ* bead method can provide better estimates for *in vivo* BSC estimation and for parameters that are derived from the BSC. In this study, we parameterized the BSC using a spherical Gaussian model to estimate the ESD and EAC. The spherical Gaussian model is a general choice to describe scattering from tissues because it is a simple model to implement and provides some geometrical interpretation of the underlying scattering structures. However, it is unknown how closely the spherical Gaussian model actually describes the underlying tissue scattering structure. While the pa-rameters have been shown to provide classification potential and reducing the bias and variance of these estimates will lead to improved classification, other models might be employed that could provide better classification. In applications that require longitudinal scans, such as scanning patients dur-ing the course of chemotherapy, the use of an *in situ* target would provide better BSC-based estimates and the ability to determine whether changes are occurring in the BSC-based parameters due to the therapy. Without an *in situ* bead target, differences in scanning orientation from one scan to the next and differences in tissue layers could result in biased estimates reducing the ability to observe changes in BSC properties due to therapy response.

## Conclusions

An *in situ* calibration approach using a 2-mm diameter Ti bead was evaluated *in vivo* in 21 tumors. The ability of the *in situ* bead method to account for attenuation and transmission losses were evaluated by comparing the BSC and BSC-based scatterer property estimates from tumors with and without slabs of lossy pork belly placed on top. The *in situ* bead method was observed to provide much lower bias on estimates when compared to the reference phantom method.

## Acknowledgements

This work was supported in part by grants from National Institutes of Health (NIH) (R21EB024133, R21EB030743, R01CA251939, R01CA273700)

## Data availability statement

The datasets generated and analyzed during the current study are accessi-ble via the following Box link: https://uofi.box.com/s/81qlbzhouzxc0pv7g1zt6fla5bsmq5w1. These datasets are publicly accessible and can be used under the terms and conditions specified at the link provided. Should there be any issues with access, please contact for assistance.

